# Vitamin K2 Limits Ferroptosis-Associated Lipid Peroxidation and Attenuates Aortic Valve Stenosis

**DOI:** 10.64898/2026.07.16.739045

**Authors:** Elena Repges, Stephan Schinhammer, Baravan Al-Kassou, Alan Yousif, Axel Schott, David Wesendonk, Benedikt Bartsch, Raul Nicolas Jamin, Malwin Barthen, Jasmin Shamekhi, Farhad Bakhtiary, Stephan Baldus, Malte Kelm, Johannes Oldenburg, Katrin J. Czogalla-Nitsche, Georg Nickenig, Sebastian Zimmer, Muntadher Al Zaidi

## Abstract

**Background:** Calcific aortic valve stenosis (AS) is the most common valvular heart disease in the aging population and lacks effective pharmacological therapy. Oxidative stress is a key feature of valvular remodeling, yet the mechanisms linking oxidative injury to calcification remain unclear. Ferroptosis, a lipid peroxidation-driven form of regulated cell death, has emerged as a key mediator of oxidative tissue injury and may contribute to cardiovascular disease. Vitamin K was recently identified as a suppressor of ferroptosis and cardiovascular calcification, but whether ferroptosis links vitamin K status to disease progression in AS remains unknown.

**Methods and Results:** We investigated the role of lipid peroxidation and ferroptosis in AS and their modulation by vitamin K2 using a translational approach. In human stenotic aortic valves, lipid peroxidation was markedly increased and localized to calcified regions, consistent with a ferroptosis-associated microenvironment. In primary human valvular interstitial cells (VICs), pro-calcific conditions induced lipid peroxidation and a pro-ferroptotic state. Pharmacological induction of ferroptosis enhanced VIC calcification, whereas its inhibition attenuated mineralization, supporting a causal role in osteogenic remodeling. Impaired vitamin K status was associated with increased valvular lipid peroxidation in AS patients. Conversely, vitamin K2 attenuated lipid peroxidation, preserved cell viability under ferroptotic stress, and partially normalized pro-ferroptotic and inflammatory transcriptional programs in VICs. In a murine model of AS, dietary vitamin K2 supplementation attenuated disease progression, reduced transvalvular gradients, and decreased valvular inflammation and lipid peroxidation-associated pathways. Finally, in a prospective cohort of patients with aortic sclerosis to moderate AS (n = 157), circulating undercarboxylated osteocalcin, a marker of impaired vitamin K status, was independently associated with accelerated disease progression.

**Conclusions:** Vitamin K2 counteracts ferroptosis-associated lipid peroxidation in AS and attenuates disease severity in vivo. Impaired vitamin K status is independently associated with accelerated progression in patients. These findings position vitamin K2 as a potential disease-modifying strategy and vitamin K status as a prognostic marker in AS.

**What Is New?:**

- Ferroptotic lipid peroxidation is enriched in calcified regions of human stenotic aortic valves. Pharmacological induction of ferroptosis increases, and its inhibition reduces, calcification of human valvular interstitial cells, indicating a causal contribution to valvular mineralization.
- Vitamin K2 suppresses ferroptotic lipid peroxidation, preserves cell viability under ferroptotic stress, and shifts pro-oxidative and pro-inflammatory transcriptional programs in human valvular interstitial cells toward a protective state.
- In an in vivo model of aortic stenosis, dietary vitamin K2 attenuated hemodynamic progression, with lower peak transvalvular velocity and mean gradient, and reduced valvular inflammation and lipid peroxidation, without affecting coagulation
- In a prospective cohort of 157 patients with aortic sclerosis to moderate stenosis, impaired vitamin K status, reflected by higher circulating undercarboxylated osteocalcin, was independently associated with accelerated disease progression and with higher rates of mortality.

**What Are the Clinical Implications?:**

- Aortic stenosis currently has no medical therapy. Vitamin K2, an inexpensive, safe nutrient that does not interfere with anticoagulation, emerges as a candidate disease-modifying strategy that warrants testing in randomized trials.
- Circulating undercarboxylated osteocalcin may serve as a biomarker to identify patients at risk of rapid progression and to enrich future vitamin K trials for those most likely to benefit.
- Targeting valvular lipid peroxidation, through antioxidant repletion and/or inhibition of lipid-peroxidation enzymes, may be a mechanistically grounded approach to slow calcific aortic valve disease.

## Introduction

Aortic valve stenosis (AS) is the most prevalent valvular heart disease in the aging population and represents a growing global health burden. Its prevalence increases steeply with age, affecting approximately 3-4% of individuals over 75 years and more than 10% of those over 80. ^1,2^ The disease is characterized by progressive fibro-calcific remodeling of the valve cusps, leading to narrowing of the aortic orifice and symptoms such as exertional dyspnea, angina, or syncope^3^. Once symptomatic, untreated AS carries a mortality of up to 50% within four years^4^. Despite its prevalence, no pharmacological therapy is currently available to halt disease progression^5,6^. A deeper understanding of the molecular mechanisms driving valvular calcification is therefore essential to identify novel therapeutic strategies.

Accumulating evidence indicates that valvular calcification is not a passive degenerative process but rather an actively regulated form of tissue remodeling. Under the influence of oxidative stress, inflammatory signaling, and lipid accumulation, valvular interstitial cells (VICs), the predominant resident cell type within the aortic valve, can undergo osteogenic differentiation and drive mineralization. However, the specific mechanisms linking oxidative stress to valvular calcification remain incompletely understood^7^.

Ferroptosis is a form of regulated cell death characterized by iron-dependent lipid peroxidation and has emerged as a key mechanism of oxidative tissue injury in degenerative disease. The process is driven by the accumulation of oxidized phospholipids, leading to membrane damage and amplification of inflammatory signaling. Ferroptosis has been implicated in several cardiovascular conditions, including atherosclerosis and heart failure^8–12^. A role in calcific aortic valve disease has recently been described, in which lipoxygenase- and ACSL4-dependent lipid peroxidation promotes valvular calcification^13^. Whether this process can be therapeutically modulated, and whether systemic vitamin K status, an endogenous determinant of lipid redox defense, relates to disease progression in patients, remains undefined.

Vitamin K–dependent pathways provide a biologically plausible link between lipid redox homeostasis and cardiovascular calcification^14^. Beyond its role in coagulation, vitamin K2 (menaquinone), the predominant extrahepatic form of vitamin K, serves as an essential cofactor for the γ-carboxylation of several proteins, including matrix Gla protein (MGP), a potent inhibitor of ectopic calcification^15^. Impaired vitamin K status, reflected by increased circulating levels of inactive vitamin K–dependent proteins, has been associated with adverse cardiovascular outcomes^16,17^. Consistently, clinical observations suggest that vitamin K antagonists accelerate valvular disease progression^18,19^, whereas dietary studies have linked higher vitamin K intake to a lower incidence of AS^20,21^. Randomized trials have yielded mixed results, with both beneficial and neutral effects reported^22,23^.

Importantly, vitamin K2 has been recently shown to modulate cellular lipid redox homeostasis and suppress ferroptosis through antioxidant mechanisms^24,25^. Whether vitamin K2 can counteract ferroptotic lipid peroxidation in the aortic valve, however, and whether impaired vitamin K status marks patients at risk of accelerated disease, has not been examined.

In the present study, we examined whether vitamin K2 modulates ferroptotic lipid peroxidation in AS and whether vitamin K status relates to disease progression. We find that lipid peroxidation is enriched in calcified regions of human aortic valves and that ferroptosis promotes VIC calcification. Extending these observations, we show that vitamin K2 attenuates lipid peroxidation and ferroptosis-associated signaling, reduces valvular remodeling in vivo, and that impaired vitamin K status, reflected by circulating undercarboxylated osteocalcin, is independently associated with disease progression in patients. These findings identify vitamin K2 as a modulator of the ferroptotic lipid-peroxidation axis in AS and establish vitamin K status as a clinical correlate of disease progression.

## Results

### Lipid Peroxidation Is Enriched in Calcified Human Aortic Valves

To determine whether lipid peroxidation is a feature of human aortic valve disease, we analyzed explanted valves from consecutive patients undergoing valve surgery for severe AS or aortic valve regurgitation associated with aortic aneurysm. Patients with active cancer or inflammatory disease were excluded, and baseline characteristics are provided in Table S1. Calcified regions from stenotic valves were compared with fibrotic and non-calcific control valve tissue from patients with aortic regurgitation. No patient was taking oral vitamin K antagonists.

Histological assessment by hematoxylin and eosin staining confirmed pronounced structural remodeling in stenotic valves, including leaflet thickening and extracellular matrix disorganization (**Figure 1A**). Consistent with advanced disease, Alizarin Red staining revealed extensive calcium deposition in stenotic valves, whereas control and fibrotic valves exhibited minimal calcification (**Figure 1B**). Quantification using OsteoSense680 demonstrated a significant increase in hydroxyapatite-positive areas in calcified valves (**Figure 1C**).

**Figure 1.**
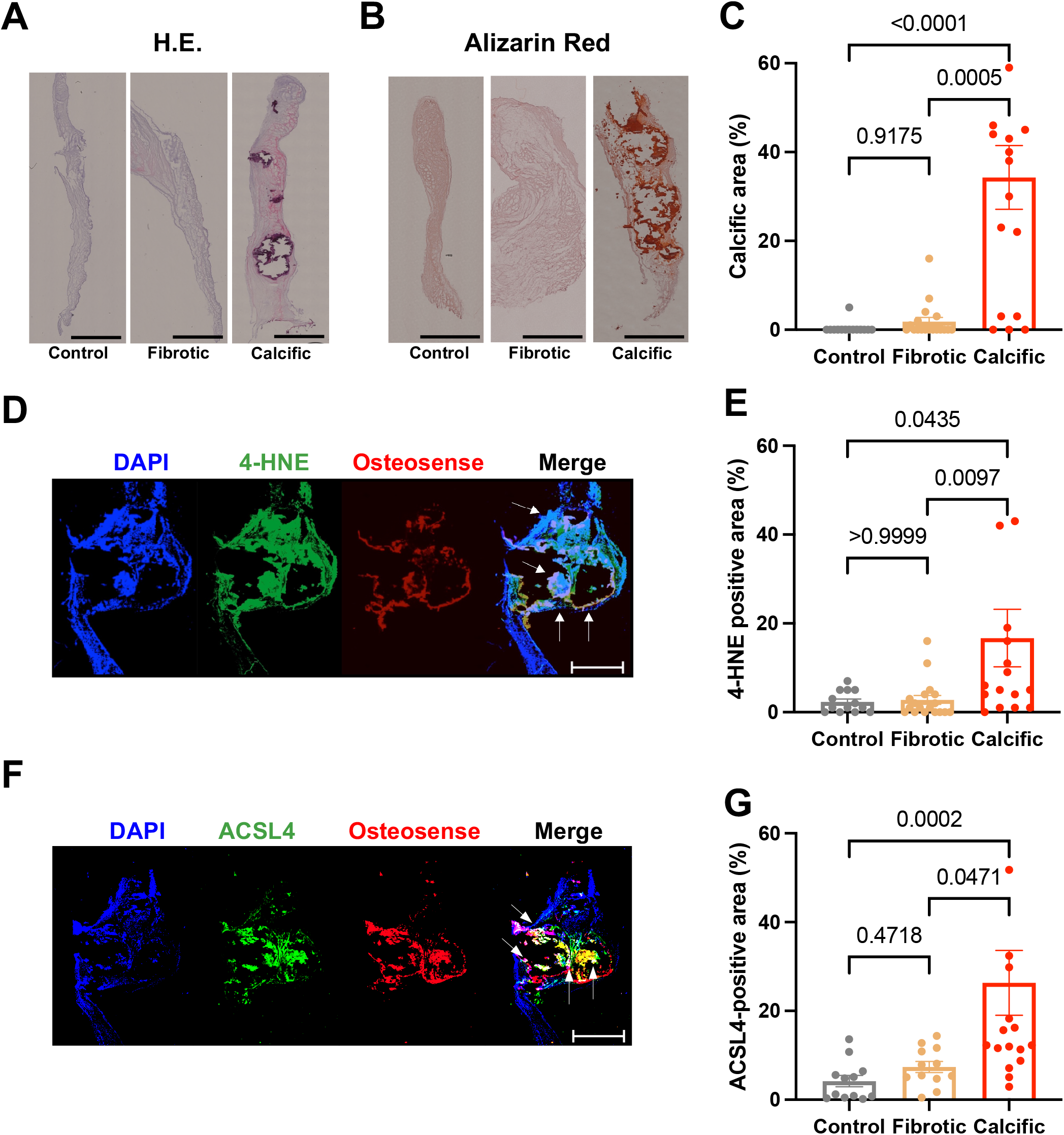
Lipid Peroxidation Is Enriched in Calcified Human Aortic Valves. (A) Representative hematoxylin and eosin staining of human aortic valves from control, fibrotic, and calcified specimens. (B) Representative Alizarin Red staining of valvular calcium deposition. (C) Quantification of calcified valve area (OsteoSense680) across conditions. (D) Representative immunostaining for 4-hydroxynonenal (4-HNE), a marker of lipid peroxidation and OsteoSense680. Arrows point to regions of overlap. (E) Quantification of 4-HNE–positive area relative to total valve area across conditions. (F) Representative immunostaining for ACSL4, a ferroptosis-associated lipid remodeling enzyme, and OsteoSense680. Arrows point to regions of overlap. (G) Quantification of ACSL4-positive area relative to total valve area across conditions. Data are presented as mean ± SEM. Statistical analysis was performed using one-way ANOVA with Tukey’s multiple-comparisons test for comparisons across valve conditions (C, E, G). Scale bars: 2 mm.

We next assessed lipid peroxidation within valve tissue. Immunostaining for 4-hydroxynonenal (4-HNE), a major end-product of lipid peroxidation, was combined with Osteosense680-staining and revealed strong overlap of pronounced lipid peroxidation signals with calcified regions (**Figure 1D**), indicating that lipid peroxidation is regionally associated with sites of mineralization. Quantification confirmed markedly increased levels in calcific valves compared with control and fibrotic tissue (**Figure 1E**). Co-localization analyses further supported spatial enrichment of 4-HNE and calcific Osteosense680-positive areas **(Figure S1)**.

To further assess lipid remodeling pathways linked to ferroptosis, we examined expression of ACSL4, a major determinant of cellular ferroptosis^26^. ACSL4 expression was similarly increased in calcified regions and co-localized with areas of mineral deposition (**Figure 1F and 1G**).

Together, these findings identify lipid peroxidation and ferroptosis-associated lipid remodeling as features of human AS that are enriched in calcified tissue.

### Ferroptosis Promotes Calcification of Human Valvular Interstitial Cells

Consistent with recent findings, and to further assess whether ferroptosis-associated pathways are enriched in human valve disease, we re-analyzed a publicly available RNA-sequencing dataset of human aortic valves comprising paired non-diseased, fibrotic, and calcific tissue from the same donors^13,27^. In a paired analysis across disease stages, calcific tissue showed increased expression of iron-import and lipoxygenase transcripts (TFRC, ALOX5, ALOX15B) together with a coordinated reduction of NRF2-linked antioxidant defenses (AKR1C1, AKR1C2, AKR1C3, SOD1) and the ferroptosis suppressor AIFM2 (**Figure 2A**; all padj < 0.05). This pattern is consistent with a shift toward a pro-ferroptotic redox state in calcified valve tissue.

**Figure 2.**
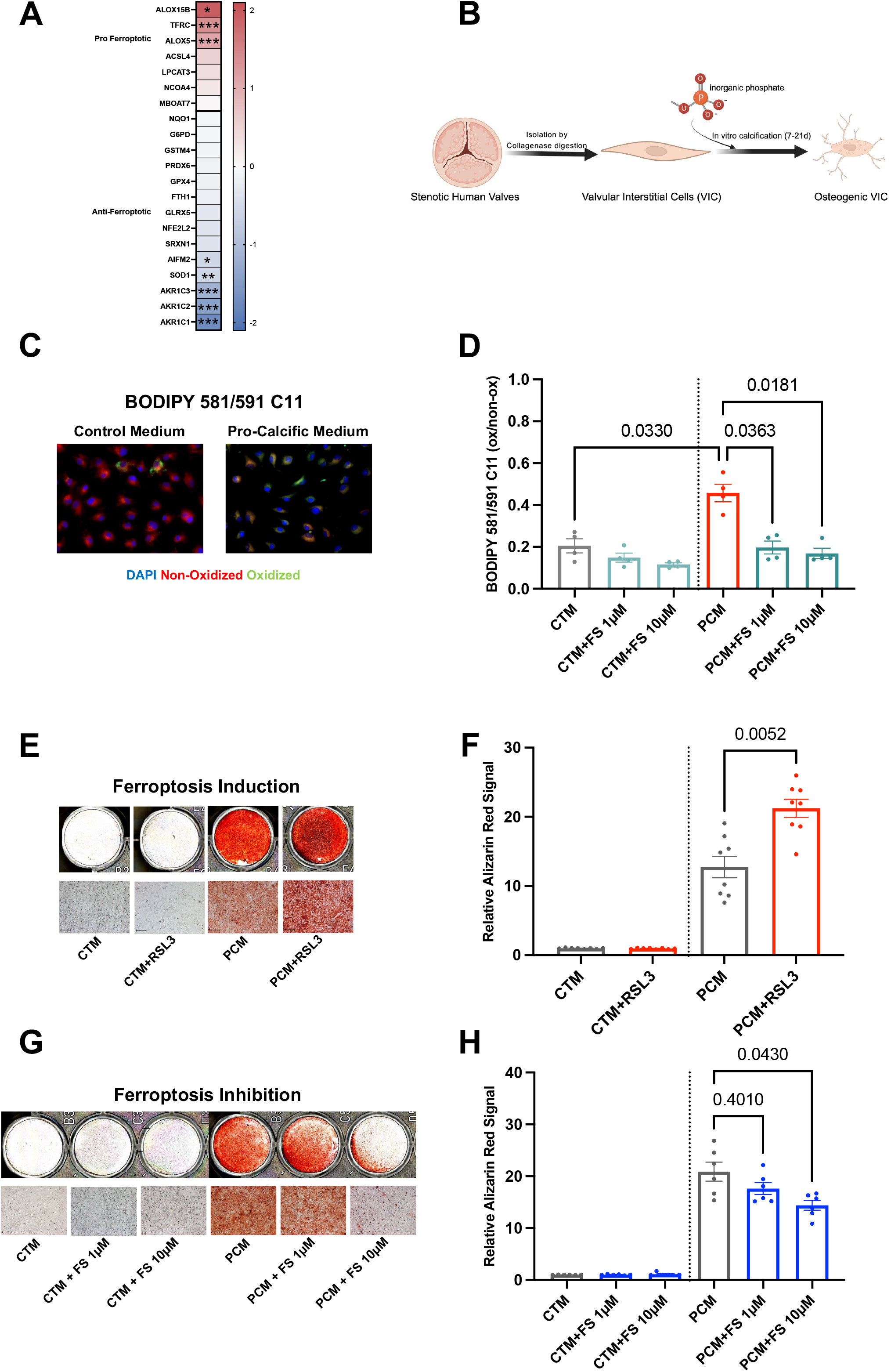
Ferroptosis Promotes Calcification of Human Valvular Interstitial Cells. (A) Ferroptosis-associated gene expression in human aortic valve tissue across disease stages (non-diseased, fibrotic, and calcific; paired samples from 3 donors). Genes are grouped as pro- or anti-ferroptotic (FerrDb-based classification). Log_2_ fold change (calcific vs. non-diseased) is shown. *padj < 0.05, **padj < 0.01, ***padj < 0.001. (B) Schematic of experimental design for valvular interstitial cells (VICs) under control and pro-calcific conditions. (C) Representative BODIPY 581/591 C11 staining illustrating lipid peroxidation in VICs (7 days of control medium or PCM). (D) Quantification of oxidized lipid species based on BODIPY fluorescence by measuring oxidized/reduced BODIPY-C11 fluorescence ratio. n = 4 independent donors. (E–F) Representative Alizarin Red staining and quantification of calcium deposition following pharmacological induction of ferroptosis with RSL3 (10nM, 21 days). n = 8 independent donors. Values were normalized to CTM. (G–H) Representative Alizarin Red staining and quantification of calcium deposition following inhibition of ferroptosis with ferrostatin-1 (1 or 10µM, 21 days). n = 6 independent donors. Values were normalized to Control Medium (CTM). Data are presented as mean ± SEM. Statistical analysis was performed using one-way ANOVA with Tukey’s multiple-comparisons test.

We next assessed, whether lipid peroxidation is induced during valvular calcification by employing a primary human valvular interstitial cell (VIC) model under pro-calcific conditions (**Figure 2B**). Exposure to pro-calcific medium (PCM) increased lipid peroxidation, as assessed by BODIPY-C11 staining (**Figure 2C**). Quantitative analysis across independent donors confirmed a shift toward oxidized lipid species under pro-calcific conditions, which was attenuated by the lipid peroxidation inhibitor ferrostatin-1 (**Figure 2D**). These findings indicate that calcifying conditions promote lipid peroxidation in VICs.

We next tested whether lipid peroxidation and ferroptotic signaling functionally contribute to VIC calcification. Pharmacological induction of ferroptosis using low-dose inhibition of GPX4 with RSL3 increased calcium deposition, as demonstrated by enhanced Alizarin Red staining and corresponding quantification (**Figure 2E–F**). Conversely, inhibition of ferroptosis with ferrostatin-1 attenuated calcification under pro-calcific conditions (**Figure 2G–H**). Collectively, these findings show that calcifying conditions induce lipid peroxidation and ferroptosis, identifying these processes as functional contributors to valvular calcification.

### Impaired Vitamin K Status Is Associated With Valvular Lipid Peroxidation and Vitamin K2 Attenuates Ferroptotic Stress in VICs

Having identified lipid peroxidation and ferroptosis-associated signaling as functional contributors to VIC mineralization, we next sought to define modulators of this process. Vitamin K emerged as a compelling candidate, given its recently described role as a potent suppressor of ferroptosis through radical-trapping antioxidant activity^24^, together with its established function in vitamin K–dependent γ-carboxylation of calcification inhibitors implicated in cardiovascular disease^15^.

We therefore first explored whether systemic vitamin K status was associated with valvular calcification and lipid peroxidation in patients with AS. Patients were stratified by median circulating undercarboxylated osteocalcin (ucOC), a marker of impaired vitamin K–dependent γ-carboxylation **(Figure 3A)**. Valves from patients with higher ucOC levels showed increased OsteoSense680-positive calcification, enhanced 4-HNE staining, and higher ACSL4 expression compared with valves from patients with lower ucOC levels **(Figure 3B–D)**. Thus, impaired systemic vitamin K status is linked to increased valvular calcification, enhanced lipid peroxidation, and ferroptosis-associated signaling in AS.

**Figure 3.**
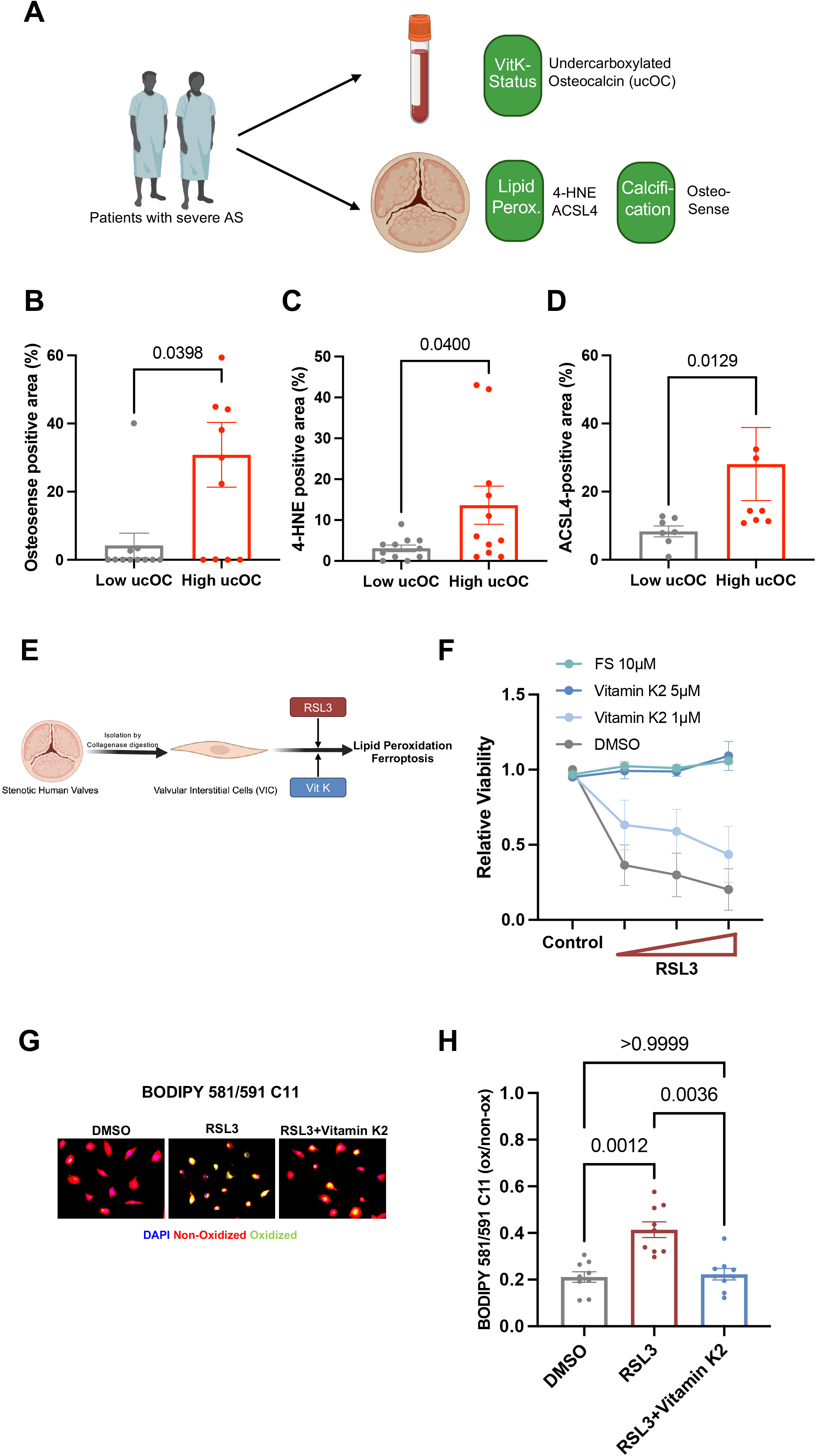
Impaired vitamin K status is associated with valvular lipid peroxidation and vitamin K2 attenuates ferroptotic stress in VICs. (A) Schematic of human sample analysis linking systemic vitamin K status (ucOC plasma levels) with valvular calcification and valvular lipid peroxidation. (B) Representative OsteoSense680 staining and quantification of calcified valve area in specimens stratified by median ucOC levels. (C) Representative 4-HNE staining and quantification of lipid peroxidation in specimens stratified by median ucOC levels. (D) Representative ACSL4 staining and quantification of ferroptosis-associated lipid remodeling in specimens stratified by median ucOC levels. (E) Schematic of experimental design for induction of ferroptosis and vitamin K2 treatment in VICs. (F) Cell viability of VICs under ferroptotic conditions following treatment with RSL3 (50nM, 100nM, 500nM, 24h), vitamin K2 (1µM or 5µM), or ferrostatin-1 (10µM). n = 3 independent donors. (G) Representative BODIPY 581/591 C11 staining illustrating lipid peroxidation following RSL3 treatment (500nM, 6h) with or without vitamin K2 (5µM). (H) Quantification of lipid peroxidation based on BODIPY fluorescence. n = 9 independent donors. Data are presented as mean ± SEM. Statistical analysis was performed using one-way ANOVA with Tukey’s multiple-comparisons test.

Based on this association between impaired vitamin K status and valvular lipid peroxidation in human tissue, we next investigated whether vitamin K2 directly modulates ferroptotic stress in primary human VICs (**Figure 3E**). To assess susceptibility to ferroptosis, VIC viability was measured following treatment with the GPX4 inhibitor RSL3. RSL3 induced a dose-dependent reduction in VIC viability, which was restored by vitamin K2 and by the ferroptosis inhibitor ferrostatin-1 (**Figure 3F)**. In parallel, RSL3 increased lipid peroxidation, as assessed by BODIPY-C11 staining (**Figure 3G)**. Co-treatment with vitamin K2 reduced lipid ROS levels compared with RSL3 alone, with quantitative analysis across independent donors confirming attenuation of lipid peroxidation (**Figure 3H**).

Together, these findings establish a link between impaired vitamin K status and valvular lipid peroxidation in human AS and demonstrate that vitamin K2 protects VICs from ferroptotic stress by reducing lipid peroxidation and preserving cell viability.

### Vitamin K2 Reprograms Calcification-, Inflammation-, and Lipid Redox–Associated Transcriptional Pathways in VICs

Having shown that vitamin K2 attenuates ferroptosis-associated lipid peroxidation under ferroptotic stress, we next asked whether vitamin K2 also modulates the broader transcriptional programs induced by calcifying conditions. To this end, we performed RNA sequencing of primary human VICs cultured under control conditions, PCM, or PCM in the presence of vitamin K2 **(Figure 4A)**.

**Figure 4.**
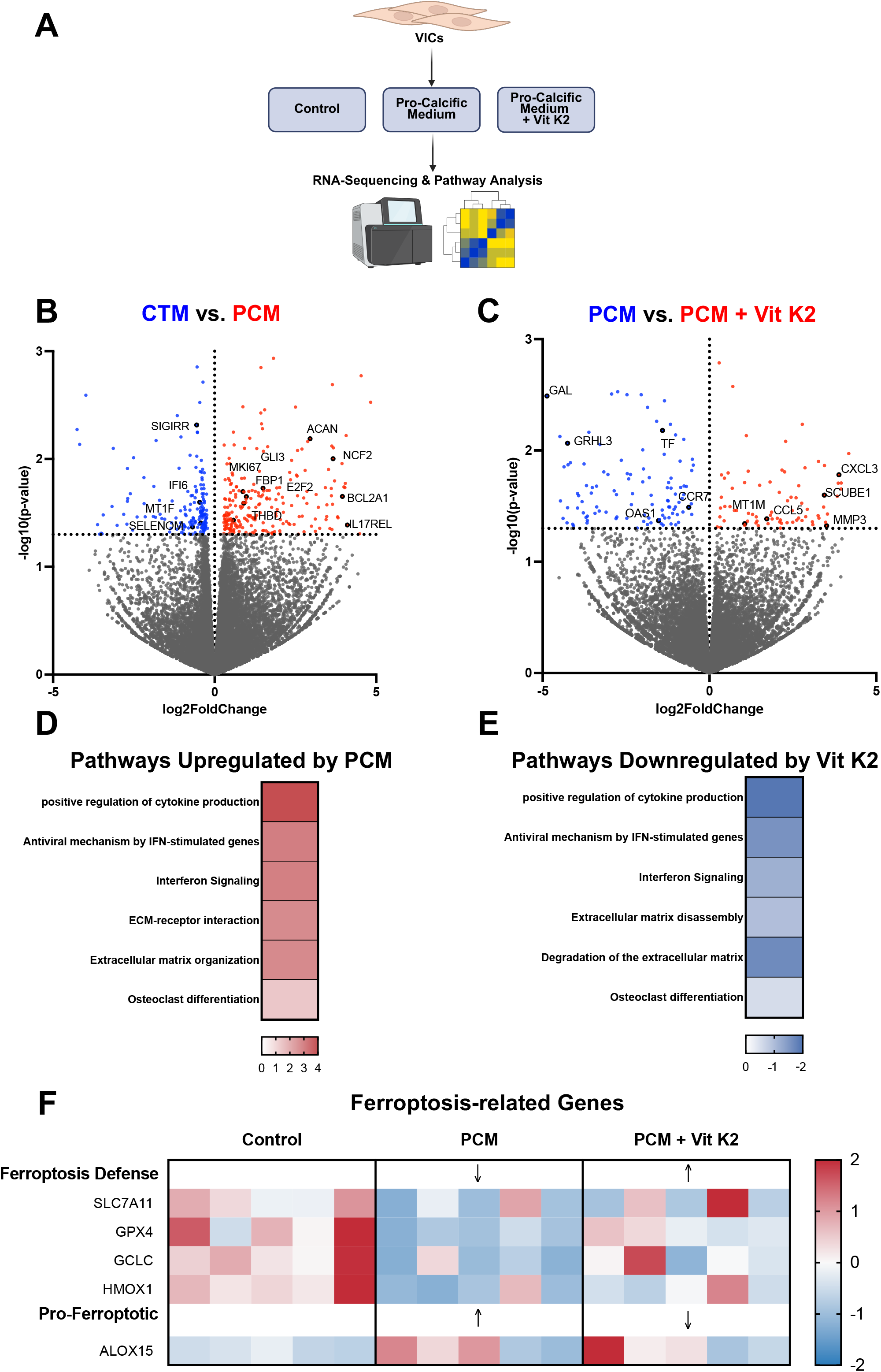
Vitamin K2 Reprograms Calcification-, Inflammation-, and Lipid Redox–Associated Transcriptional Pathways in VICs. (A) Schematic of RNA sequencing experimental design in VICs cultured under control, pro-calcific, and vitamin K2–treated conditions (n=5 independent donors). (B) Volcano plot illustrating differential gene expression in VICs under pro-calcific conditions compared with control. (C) Volcano plot illustrating differential gene expression following vitamin K2 treatment and pro-calcific medium compared with pro-calcific conditions alone. (D) Pathway enrichment analysis (GO Terms/KEGG/Reactome) of upregulated pathways under pro-calcific conditions. (E) Pathway enrichment analysis (GO Terms/KEGG/Reactome) following vitamin K2 treatment. (F) Heatmap showing normalized expression of lipid redox–related genes, including *SLC7A11, GCLC, GPX4, HMOX1,* and *ALOX15*. Differential expression was analyzed using DESeq2.

Differential gene expression analysis revealed marked transcriptional changes in VICs exposed to pro-calcific conditions compared with control medium, as illustrated by volcano plot analysis (**Figure 4B**). Vitamin K2 treatment induced a distinct transcriptional profile compared with PCM alone (**Figure 4C**), indicating modulation of calcification-associated gene expression.

Pathway enrichment analysis showed that pro-calcific conditions were associated with upregulation of pathways related to cytokine signaling, interferon responses, and extracellular matrix remodeling (**Figure 4D**). In contrast, vitamin K2 treatment attenuated these responses and was associated with enrichment of pathways linked to cellular homeostasis and stress adaptation (**Figure 4E**).

We next examined genes involved in lipid redox regulation. Pro-calcific conditions were associated with reduced expression of antioxidant and lipid repair systems, including *SLC7A11*, *GCLC*, *GPX4*, and *HMOX1*, alongside increased expression of *ALOX15*, an enzyme involved in lipid peroxidation. Vitamin K2 treatment partially reversed this pattern, restoring expression of antioxidant genes while reducing *ALOX15* expression (**Figure 4F**).

Together, these data indicate that pro-calcific conditions are associated with coordinated changes in inflammatory, extracellular matrix, and lipid redox pathways, and that vitamin K2 modulates these transcriptional programs toward a less pro-oxidative and less pro-inflammatory state.

### Vitamin K2 Attenuates AS and Modulates Inflammatory and Lipid Peroxidation–Associated Remodeling In Vivo

Based on our findings that vitamin K2 attenuates ferroptosis-associated lipid peroxidation and modulates calcification-associated transcriptional programs in VICs, we next assessed whether vitamin K2 affects AS disease progression in vivo. To this end, we used a murine wire injury–induced model of AS combined with dietary vitamin K2 supplementation **(Figure 5A)**. Vitamin K2 supplementation was initiated directly after wire injury and continued throughout follow-up.

**Figure 5.**
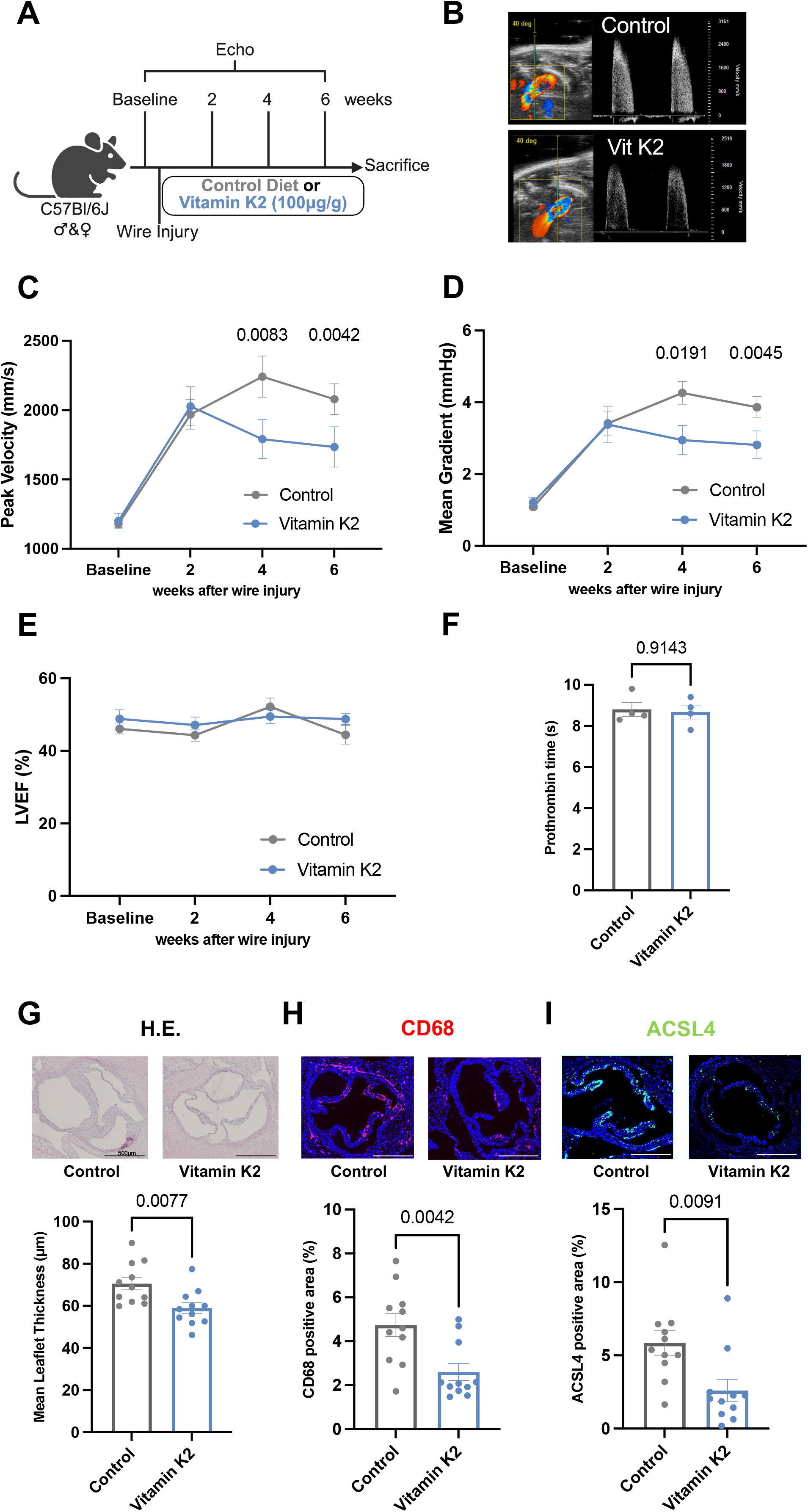
Vitamin K2 Attenuates Experimental AS and Modulates Inflammatory and Ferroptosis-associated Remodeling In Vivo. (A) Schematic of the murine wire injury–induced model of AS with dietary vitamin K2 (menaquinone-7, 100µg/g chow) supplementation. (B) Representative Doppler echocardiographic images illustrating transvalvular flow in control and vitamin K2–treated mice. (C-E) Serial assessment of peak transvalvular velocity (C), mean transvalvular gradient (D), and left ventricular ejection fraction (E) by echocardiography. (F) Assessment of systemic coagulation by plasma prothrombin time in a subset of n = 4 per group. (G) Representative hematoxylin and eosin staining of aortic valve leaflets and quantification of leaflet thickness. (H) Representative immunofluorescence staining for CD68 and quantification of valvular macrophage accumulation. (I) Representative immunofluorescence staining for ACSL4 and quantification of valvular expression. Data are presented as mean ± SEM. Statistical analysis in (C&D) was performed using 2-way ANOVA with Tukey’s multiple-comparisons test. Statistical analyses in (F–I) were performed using unpaired 2-tailed Student’s t test. n = 11 per group. Scale bar: 500µm.

Serial echocardiography demonstrated progressive increases in peak transvalvular velocity in control animals, whereas vitamin K2-treated mice exhibited significantly attenuated progression over time **(Figure 5B-C)**. Consistently, mean transvalvular gradients were lower in vitamin K2-treated animals **(Figure 5D)**, indicating reduced hemodynamic severity of stenosis. Left ventricular systolic function remained preserved in both groups, with no differences in ejection fraction **(Figure 5E)**. Importantly, vitamin K2 supplementation did not affect systemic coagulation, as assessed by prothrombin time **(Figure 5F)**.

Histological analysis revealed pronounced leaflet thickening and structural remodeling in control animals, which were attenuated by vitamin K2 treatment **(Figure 5G)**. Immunostaining for CD68 demonstrated reduced macrophage accumulation in vitamin K2-treated valves **(Figure 5H)**, consistent with lower circulating monocyte counts **(Figure S2A)**. Plasma proteomic profiling further supported systemic modulation of inflammation- and calcification-associated circulating proteins **(Figure S2B)**, whereas valvular collagen content was not significantly altered **(Figure S2C)**.

Given our clinical and in vitro findings linking ferroptosis-associated lipid peroxidation to vitamin K status in AS, we next assessed lipid peroxidation-associated pathways in vivo. Expression of ACSL4, an enzyme dictating ferroptosis sensitivity by modulating lipid remodeling processes^26^, was reduced in valves from vitamin K2–treated animals **(Figure 5I)**.

Together, these findings demonstrate that vitamin K2 attenuates experimental AS in vivo and is associated with reduced inflammatory remodeling and lower ferroptosis-associated lipid remodeling. These data provide in vivo support for the vitamin K2–sensitive lipid peroxidation axis identified in human samples and primary VICs.

### Impaired Vitamin K Status Is Associated With Accelerated Progression of Aortic Valve Disease

Finally, we assessed the clinical relevance of vitamin K status for disease progression in patients with AS. We analyzed a prospective cohort of n = 157 patients spanning the spectrum from aortic sclerosis to moderate AS, thereby enabling assessment of progression across the continuum of aortic valve disease **(Figure 6A)**. The median age was 74 years (IQR 68–81), and 63.1% of participants were male. The prevalence of cardiovascular risk factors and comorbidities included hypertension (89.2%), hypercholesterolemia (77.1%), diabetes mellitus (29.2%), coronary artery disease (42.9%), and chronic kidney disease (41.5%). At baseline, peak transvalvular velocity and mean gradient were 2.69 ± 0.50 m/s and 15.3 ± 6.4 mmHg, respectively, while left ventricular ejection fraction was preserved (59.3% [IQR 55.1–64.6]). Patients underwent standardized clinical assessment and blood sampling at study entry, followed by serial echocardiographic evaluation over a median follow-up of 776 days (IQR 399–935). No patient was taking oral vitamin K antagonists.

**Figure 6.**
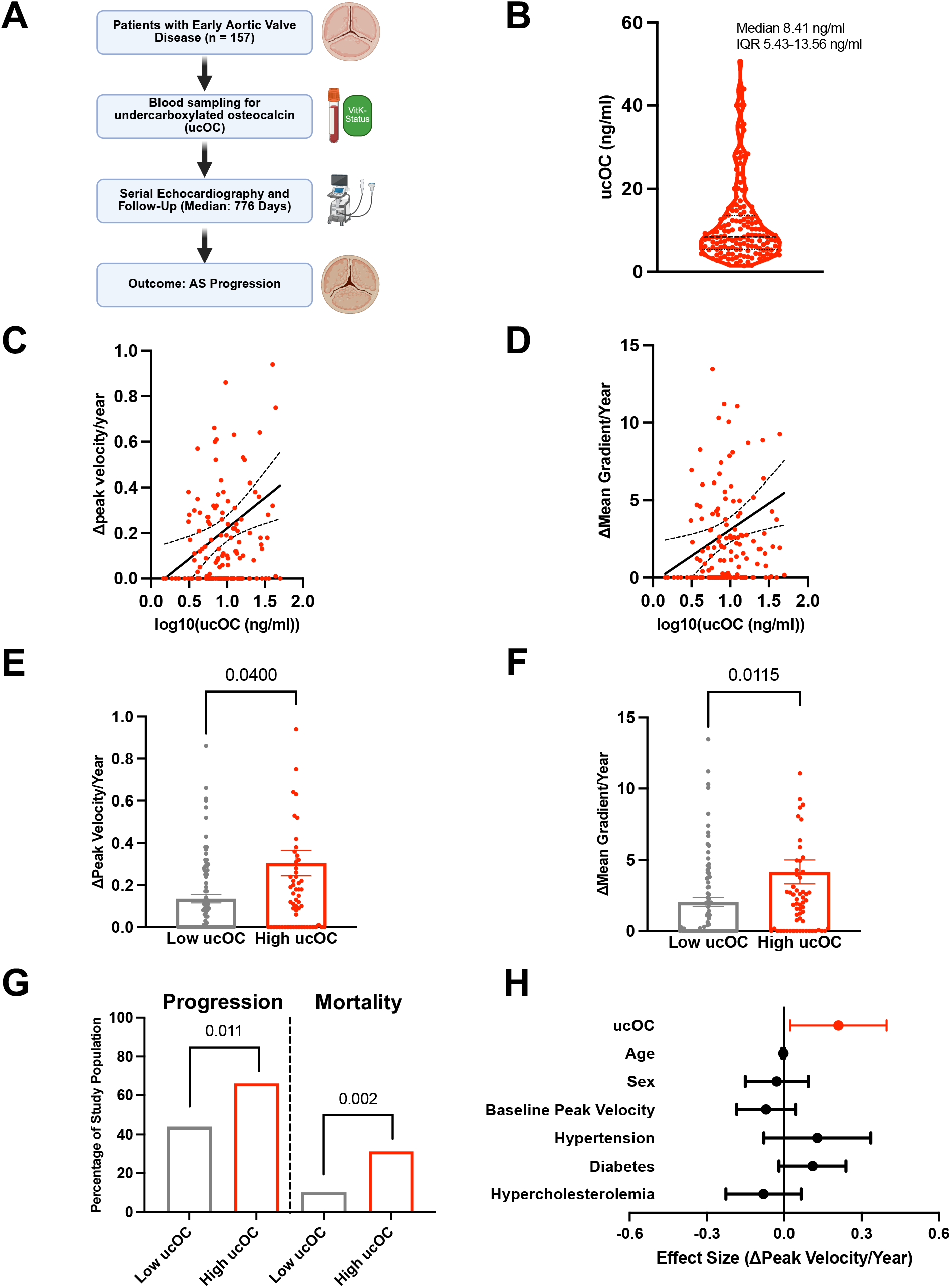
Impaired vitamin K Status Is Associated with Accelerated Progression of Aortic Valve Stenosis. (A) Study design of the prospective patient cohort (n = 157). (B) Distribution of circulating undercarboxylated osteocalcin (ucOC) levels. (C–D) Association of baseline ucOC levels with annualized progression of peak transvalvular velocity and mean transvalvular gradient. (E–F) Stratified analysis of disease progression according to ucOC levels using a Youden-derived cut-off. (G) Incidence rates of AS disease progression and mortality stratified by ucOC levels throughout follow-up. (H) Multivariable regression analysis assessing the association of ucOC with disease progression. Data are presented as mean ± SEM or median (IQR), as appropriate. Statistical analysis was performed using linear regression (C-D, H), unpaired 2-tailed Student’s t test (E-F), and chi square test (G), as appropriate.

Vitamin K status was assessed by circulating levels of undercarboxylated osteocalcin (ucOC), a marker of impaired vitamin K–dependent γ-carboxylation. Baseline ucOC levels exhibited a broad distribution within the cohort (median 8.41 ng/mL, IQR 5.43–13.56 ng/mL) **(Figure 6B)**. Baseline characteristics stratified by ucOC levels are shown in **Table 1** and were well balanced between groups except for a higher incidence of chronic kidney disease in patients with high ucOC levels.

**Table 1.**
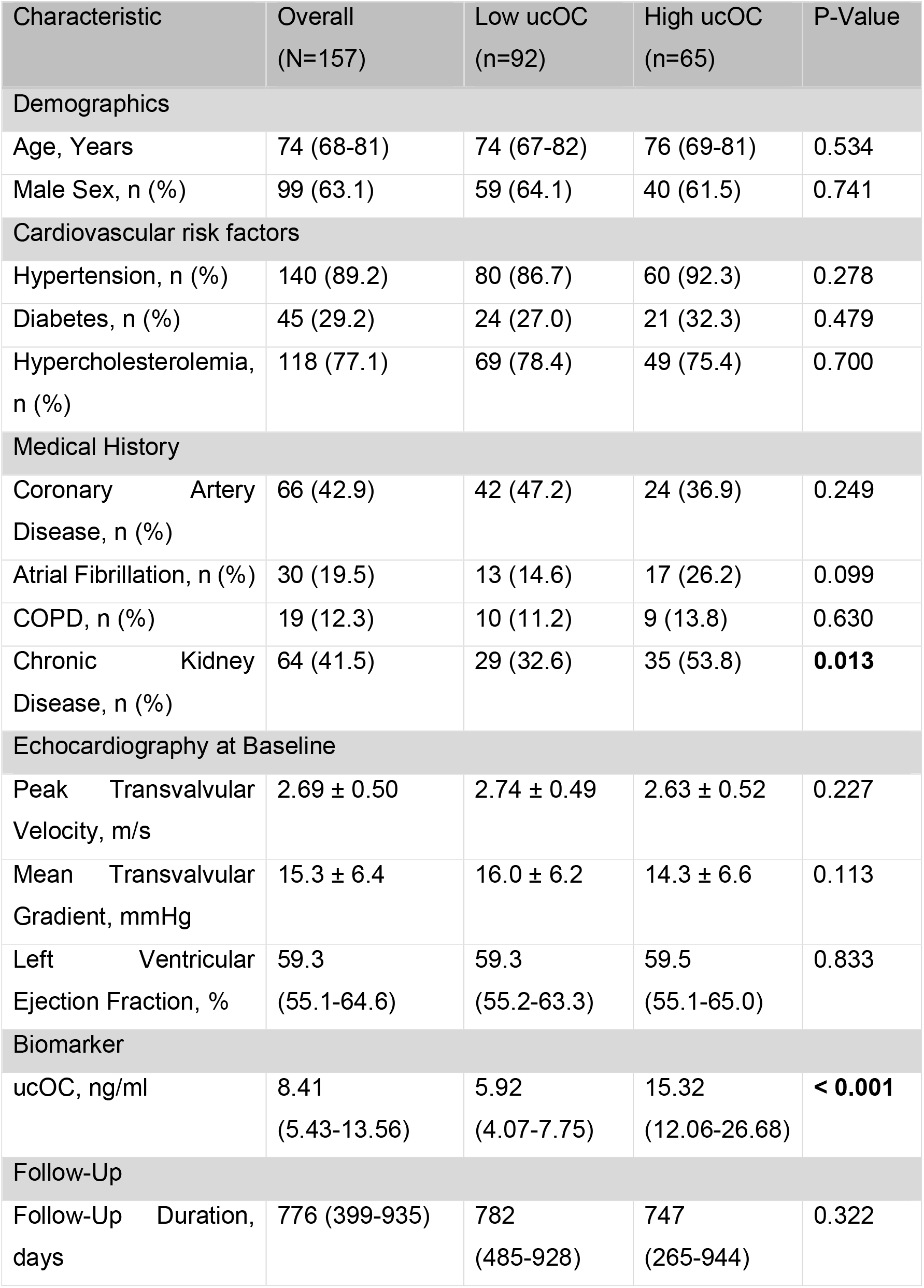
Baseline Characteristics.

| Characteristic | Overall<br>(N=157) | Low ucOC<br>(n=92) | High ucOC<br>(n=65) | P-Value |
| --- | --- | --- | --- | --- |
| Demographics |  |  |  |  |
| Age, Years | 74 (68-81) | 74 (67-82) | 76 (69-81) | 0.534 |
| Male Sex, n (%) | 99 (63.1) | 59 (64.1) | 40 (61.5) | 0.741 |
| Cardiovascular risk factors |  |  |  |  |
| Hypertension, n (%) | 140 (89.2) | 80 (86.7) | 60 (92.3) | 0.278 |
| Diabetes, n (%) | 45 (29.2) | 24 (27.0) | 21 (32.3) | 0.479 |
| Hypercholesterolemia,<br>n (%) | 118 (77.1) | 69 (78.4) | 49 (75.4) | 0.700 |
| Medical History |  |  |  |  |
| Coronary Artery<br>Disease, n (%) | 66 (42.9) | 42 (47.2) | 24 (36.9) | 0.249 |
| Atrial Fibrillation, n (%) | 30 (19.5) | 13 (14.6) | 17 (26.2) | 0.099 |
| COPD, n (%) | 19 (12.3) | 10 (11.2) | 9 (13.8) | 0.630 |
| Chronic Kidney<br>Disease, n (%) | 64 (41.5) | 29 (32.6) | 35 (53.8) | **0.013** |
| Echocardiography at Baseline |  |  |  |  |
| Peak Transvalvular<br>Velocity, m/s | 2.69 ± 0.50 | 2.74 ± 0.49 | 2.63 ± 0.52 | 0.227 |
| Mean Transvalvular<br>Gradient, mmHg | 15.3 ± 6.4 | 16.0 ± 6.2 | 14.3 ± 6.6 | 0.113 |
| Left Ventricular<br>Ejection Fraction, % | 59.3<br>(55.1-64.6) | 59.3<br>(55.2-63.3) | 59.5<br>(55.1-65.0) | 0.833 |
| Biomarker |  |  |  |  |
| ucOC, ng/ml | 8.41<br>(5.43-13.56) | 5.92<br>(4.07-7.75) | 15.32<br>(12.06-26.68) | **< 0.001** |
| Follow-Up |  |  |  |  |
| Follow-Up Duration,<br>days | 776 (399-935) | 782<br>(485-928) | 747<br>(265-944) | 0.322 |

Higher baseline ucOC levels were associated with accelerated progression of AS. In continuous analyses, ucOC was significantly associated with a greater annualized increase in both peak transvalvular velocity (β = 0.27 per log10 increase in ucOC, 95% CI 0.09–0.45; p = 0.004) and mean transvalvular gradient (β = 3.39, 95% CI 0.82–5.95; p = 0.010) **(Figure 6C and 6D)**. Stratification by ucOC levels (Youden-derived cut-off: 9.73 ng/mL) demonstrated that patients with elevated ucOC exhibited significantly faster disease progression compared with those with lower levels (Δpeak velocity: 0.11 vs 0.05 m/s/year, p = 0.040; Δmean gradient: 2.00 vs 0.39 mmHg/year, p = 0.0115) **(Figure 6E and 6F)**. Consistent with these findings, binary stratification showed that patients with higher ucOC levels had significantly higher rates of both disease progression (increase of at least one stage in AS grading) and mortality compared with those with lower ucOC levels **(Figure 6G)**.

In multivariable regression analyses adjusting for age, sex, baseline peak velocity, and cardiovascular risk factors including hypertension, diabetes mellitus, and hypercholesterolemia, ucOC remained independently associated with accelerated AS progression (adjusted β = 0.21, 95% CI 0.02–0.39; p = 0.028) **(Figure 6H)**. Given the association between ucOC and chronic kidney disease, additional sensitivity analyses were performed in patients without CKD, in whom ucOC remained independently associated with disease progression (adjusted β = 0.20, 95% CI 0.01–0.39; p = 0.039).

Together, these findings extend our experimental observations to a longitudinal patient cohort and identify impaired vitamin K status as an independent predictor of AS progression.

## Discussion

In this study, we show that vitamin K2 counteracts ferroptotic lipid peroxidation in AS and that impaired vitamin K status is associated with disease progression. Across human valve tissue, primary VICs, experimental AS, and a prospective clinical cohort, our data confirm that ferroptotic signaling contributes to valvular calcification and, extending this concept, identify vitamin K2 as a modulator of this axis and vitamin K status as a clinical correlate of progression.

A central observation is the marked enrichment of lipid peroxidation within calcified regions of human stenotic valves. AS is increasingly recognized as an active, multicellular disease process initiated by endothelial dysfunction, lipid infiltration, and immune cell recruitment, ultimately driving VIC activation and osteogenic differentiation^7,28^. While oxidative stress has long been implicated in this cascade^29^, our data refine this concept by identifying lipid peroxidation as a spatially localized form of oxidative injury within diseased valves. In the context of lipid accumulation and redox-active iron, the valvular microenvironment appears permissive for ferroptotic signaling^30–33^. These findings align with growing evidence that ferroptosis contributes to AS and other chronic cardiovascular diseases^13^, including atherosclerosis^8,12^, heart failure^10^, and aortic aneurysm^11^.

Mechanistically, lipid peroxidation is not merely a byproduct of tissue injury but directly contributes to VIC dysfunction and calcification. Ferroptotic processes involve iron-dependent peroxidation of polyunsaturated phospholipids, leading to accumulation of reactive lipid species such as 4-hydroxynonenal, which disrupt membrane integrity and amplify inflammatory signaling^34,35^. Importantly, lipid peroxidation products have been shown to promote osteogenic differentiation, for example through modification of RUNX2, supporting a causal link between oxidative lipid damage and mineralization^36^. These findings position lipid peroxidation as a key mediator linking cellular stress, inflammation, and calcific remodeling.

Our transcriptomic analyses support this model, demonstrating that pro-calcific conditions induce coordinated activation of inflammatory, extracellular matrix, and lipid redox pathways. Ferroptosis and inflammation are tightly interconnected processes^37^: lipid peroxidation products amplify cytokine signaling and immune cell recruitment^38–40^, while inflammatory mediators further promote phospholipid remodeling and oxidative injury^41,42^. Together, these data indicate that lipid peroxidation operates within a self-reinforcing network that drives valvular remodeling.

Our data further define vitamin K2 as a modulator of this axis. In addition to its established role in γ-carboxylation of proteins such as matrix Gla protein ^15,19,43^, vitamin K2 acts as a lipophilic radical-trapping antioxidant that limits ferroptotic lipid peroxidation^24,44^. This activity depends on intracellular recycling by enzymes such as FSP1^24^ and the VKORC1-isoenzyme VKORC1L1, which is highly expressed in extrahepatic tissues and sustains cellular antioxidant defense^25,45–47^. These findings are consistent with earlier biochemical observations demonstrating antioxidant properties of vitamin K2 within lipid environments^48^. In the present study, vitamin K2 reduced lipid peroxidation, improved cellular resilience under ferroptotic stress, and partially reversed pro-ferroptotic transcriptional programs. Consistent with these findings, dietary vitamin K2 supplementation attenuated the progression of experimental AS and reduced inflammatory remodeling in vivo. These observations suggest that modulation of lipid peroxidation influences both VIC-intrinsic stress responses and the inflammatory microenvironment.

Clinical data extend these findings. In a prospective cohort, impaired vitamin K status was independently associated with accelerated disease progression. In accordance, prior observational studies have linked low vitamin K status to increased cardiovascular calcification and AS risk^20,21^, while pharmacological inhibition of the vitamin K cycle by vitamin K antagonists has been associated with accelerated valvular calcification^18,19,49^. Interventional studies, however, have yielded mixed results. A randomized trial in patients with mild-to-moderate AS demonstrated slower calcification progression with vitamin K supplementation^22^, whereas another trial in individuals with early-stage valve calcification showed no effect on disease progression^23^.

These differences likely reflect variation in disease stage, patient selection, baseline vitamin K status, and concomitant treatments. Our findings raise the possibility that vitamin K2 may be most effective in patient subsets characterized by active lipid peroxidation and ferroptotic signaling, suggesting that using biomarkers of vitamin K deficiency or lipid peroxidation could improve patient stratification in future trials.

Several limitations should be acknowledged. The observational nature of the clinical data precludes causal inference. Experimental models, while providing mechanistic insight, do not fully recapitulate the complexity of human disease. In addition, the present study does not fully distinguish γ-carboxylation-dependent from non-canonical antioxidant effects of vitamin K2. Finally, the specific lipid species driving VIC dysfunction and calcification remain to be defined and warrant further investigation.

In conclusion, vitamin K2 counteracts ferroptosis-associated lipid peroxidation in AS and attenuates experimental disease, and impaired vitamin K status is independently associated with accelerated progression. Targeting lipid peroxidation with vitamin K2 may represent a disease-modifying strategy in AS, and vitamin K status may help identify patients most likely to benefit.

## Methods

### Human Aortic Valve Tissue

Human aortic valve tissue and blood samples were obtained from patients undergoing surgical valve replacement for severe AS or from control patients undergoing surgery for aortic regurgitation at the Heart Center Bonn, Germany. All patients provided written informed consent, and the study was approved by the Ethics Committee of the University of Bonn (approval number 078/17). Immediately after excision, valve cusps were rinsed in ice-cold phosphate-buffered saline (PBS).

For regional analyses, valve tissue was macroscopically dissected into calcified, fibrotic, and non-diseased areas. Samples were either embedded in Tissue-Tek freezing medium for histological analysis or snap-frozen in liquid nitrogen and stored at −80°C until further use.

Cryosections were air-dried at room temperature and processed for histological and immunofluorescence analyses. Staining was performed as described below in the respective section. For morphological assessment, sections were stained with hematoxylin and eosin and calcification was assessed using Alizarin Red staining using standard protocols. For Osteosense680-staining, slides were incubated with the Osteosense680 probe (1:100 in 2% BSA in PBS, PerkinElmer #NEV10020EX) for 24 hours at 4°C in the dark. Slides were then washed four times with PBS and immunostaining with primary and secondary antibodies was performed as described below.

Images were acquired using a fluorescence microscope (Zeiss Axio Observer or equivalent) under identical exposure settings across groups. Multiple regions per valve were imaged. Quantification of lipid peroxidation, ACSL4 expression, and calcified areas was performed using ImageJ by applying consistent thresholding across all samples. Signals were expressed relative to total tissue area. All analyses were performed by investigators blinded to grouping.

### Isolation and Culture of Valvular Interstitial Cells

Primary human valvular interstitial cells (VICs) were isolated from aortic valve cusps by enzymatic digestion. Tissue was minced and incubated in collagenase II (600 U/mL; Thermo Fisher Scientific, 21885108) in Dulbecco’s Modified Eagle Medium at 37°C with gentle agitation for 20 hours. The resulting cell suspension was filtered through a 100 µm cell strainer, centrifuged at 1000 rpm for 15 minutes, and resuspended in VIC culture medium consisting of Dulbecco’s Modified Eagle Medium supplemented with 10% fetal bovine serum and 1% penicillin/streptomycin. VICs were cultured at 37°C in a humidified atmosphere containing 5% CO₂. Medium was changed every 2 to 3 days. Experiments were performed using cells between passages 4 and 6.

### In Vitro Calcification Model

VICs were seeded in 12-well plates and cultured in control medium or pro-calcific medium (PCM). PCM consisted of Dulbecco’s Modified Eagle Medium supplemented with 5% fetal bovine serum, 1% penicillin/streptomycin, 2 mmol/L sodium dihydrogen phosphate, and 50 µg/mL L-ascorbic acid. VICs were stimulated for up to 21 days for assessment of calcification and for 24 to 72 hours for molecular and lipid peroxidation analyses. Medium was replaced every 2 to 3 days.

### Pharmacological Modulation of Ferroptosis

Ferroptosis was induced using the GPX4 inhibitor RSL3 (10 - 500 nM; MedChemExpress, #HY-100218A) for 24 hours to 21 days for calcification experiments. Ferroptosis was inhibited using ferrostatin-1 (1 - 10 µM; MedChemExpress, HY-100579). Vitamin K2 (menaquinone-7, MK-7, a generous gift of Gnosis by Lesaffre International S.A.S, Marcq-en-Baroeul, France) was dissolved in dimethyl sulfoxide and applied at concentrations of 1 - 5 µM. Control conditions contained equivalent concentrations of vehicle (DMSO).

### Alizarin Red Staining and Quantification

After 21 days of stimulation under control or pro-calcific conditions, VICs were fixed in 4% paraformaldehyde for 15 minutes at room temperature and washed with distilled water. Cells were stained with 2% Alizarin Red S solution (pH 4.2) for 15 minutes and washed extensively with distilled water. For quantification, bound dye was extracted using 10% cetylpyridinium chloride (dissolved in ddH_2_O) for 1 hour at room temperature. Absorbance was measured at 550 nm using a microplate reader.

### Lipid Peroxidation Assay

Lipid peroxidation in VICs was assessed using BODIPY 581/591 C11 (Thermo Fisher Scientific). Cells were incubated with 5 µM dye for 20 minutes at 37°C protected from light. Nuclei were stained with Hoechst 33342 (1 µg/ml, Thermo Fisher # 62249) for five minutes. After washing, VICs were fixed for fluorescence microscopy. Oxidized and reduced lipid signals were detected in the FITC and PE channels and relative oxidation levels were quantified using ImageJ.

### Cell Viability Assay

Cell viability of VICs under ferroptotic stress conditions was assessed using alamarblue HS cell viability assay (Thermo Fisher, # A50101) according to the manufacturer’s instructions. Absorbance was measured at 570 nm with wavelength correction at 600 nm using a microplate reader (Infinite M200, Tecan).

### RNA Isolation

Total RNA was extracted from VICs using TRIzol reagent according to the manufacturer’s protocol. RNA concentration was determined using a NanoDrop spectrophotometer.

### RNA Sequencing and Bioinformatic Analysis

RNA sequencing was performed by Novogene (Cambridge, UK). RNA integrity was assessed using an Agilent 5400 Bioanalyzer, and only samples with RNA integrity number (RIN) ≥9 were used. Sequencing libraries were generated from 200 ng total RNA using the QuantSeq 3′ mRNA-Seq Library Prep Kit (Lexogen) according to the manufacturer’s instructions, including poly(A) selection for mRNA enrichment. Libraries were indexed, pooled, and sequenced on an Illumina NovaSeq X Plus platform. Raw sequencing reads were subjected to quality control using FastQC, and adapter sequences and low-quality reads were removed. Differential gene expression analysis was performed in R using DESeq2. P values were adjusted for multiple testing using the Benjamini–Hochberg method.

### Analysis of Public Transcriptomic Data

Publicly available RNA-sequencing data of human aortic valves (non-diseased, fibrotic, and calcific tissue from 3 paired donors) were re-analyzed^27^. Raw counts were filtered (total count ≥ 10) and analyzed with DESeq2 using a paired design (donor + stage). Differential expression between disease stages was tested by Wald test with Benjamini-Hochberg correction. A curated ferroptosis gene panel (drivers and suppressors, based on FerrDb and primary literature) was visualized as a heatmap.

### Animal Experiments

Male and female C57BL/6J mice aged 10–12 weeks (Janvier Labs) were used. Animals were housed under specific pathogen–free conditions with a 12-hour light/dark cycle and ad libitum access to food and water. All experiments were approved by the respective authority (LANUV/LAVE North Rhine-Westphalia; file number 81-02.04.2023.A390) and performed in accordance with institutional guidelines and Directive 2010/63/EU of the European Parliament. Experiments were conducted and reported in accordance with ARRIVE guidelines.

Mice were randomly assigned to receive either standard chow or chow supplemented with vitamin K2 (menaquinone-7: MK-7, a generous gift of Gnosis by Lesaffre International S.A.S, Marcq-en-Baroeul, France, 100 µg/g diet; ssniff-Spezialdiäten GmbH, Soest, Germany). Allocation was performed using a randomized block design with balanced distribution by sex. Investigators performing surgical procedures, echocardiographic assessment, and data analysis were blinded to group allocation. Sample size was determined based on prior experience with the model and expected effect sizes. No animals were excluded from analysis unless predefined humane endpoints were reached.

### Wire Injury Model

Aortic valve injury was induced as previously described^50,51^. Mice were anesthetized with intraperitoneal ketamine (100 mg/kg) and xylazine (16 mg/kg). Adequate anesthesia was confirmed by loss of pedal reflex. Body temperature was maintained at 37°C using a heating pad. Following a right paramedian cervical incision, the right common carotid artery was exposed. A coronary guide wire with a 15° angled tip was introduced into the artery and advanced retrogradely across the aortic valve under echocardiographic guidance. Valve injury was induced by standardized mechanical manipulation consisting of 100 rotations and 10 longitudinal passages across the valve. After the procedure, the carotid artery was ligated, and the incision was closed in layers. Postoperative analgesia was provided using buprenorphine. Animals were monitored daily for recovery and signs of distress.

### Echocardiography

Transthoracic echocardiography was performed at baseline and at predefined time points using a high-frequency ultrasound system (Vevo 2100, Fujifilm VisualSonics). Mice were anesthetized with 1.5% isoflurane in oxygen, and heart rate, respiratory rate, and body temperature were continuously monitored. Aortic valve peak transvalvular velocity was assessed using pulse-wave Doppler in the suprasternal view with angle correction (40°). All echocardiographic analyses were performed by investigators blinded to treatment allocation. Measurements were averaged from at least three consecutive cardiac cycles.

### Prothrombin Time

Prothrombin time was measured in 50µl of citrated plasma using Innovin^®^ Quick reagent assay according to the manufacturer’s protocol using a MC10 device.

### Histological Analysis of Murine Valves

For morphological assessment, cryosections were air-dried at room temperature and stained with hematoxylin and eosin according to standard procedures. Leaflet thickness was quantified on H&E-stained sections by measuring maximal cusp thickness at nine predefined anatomical landmarks using ImageJ (NIH). Measurements were averaged from at least three sections per animal.

### Immunofluorescence

For immunofluorescence, cryosections were fixed in freshly prepared cold acetone for 20 minutes at room temperature, followed by four washes in phosphate-buffered saline (PBS). Sections were encircled with a hydrophobic barrier and blocked with 2% bovine serum albumin (BSA) in PBS for 30 minutes at room temperature.

Primary antibody incubation was performed overnight at 4°C using antibodies against CD68 (1:100 in 2% BSA; antibodies-online, #ABIN181836), ACSL4 (1:50 in 2% BSA, Abcam #ab155282), 4-HNE (1:100 in 2% BSA, Abcam #ab46545). After washing four times with PBS (5 minutes each), sections were incubated with species-appropriate fluorophore-conjugated secondary antibodies (Cy3, 1:500 in 2% BSA; Jackson ImmunoResearch # 712-165-153 or Goat IgG anti-Rabbit IgG (H+L)-Cy3, Dianova 111-165-144) for 1 hour at room temperature in the dark. Sections were washed again in PBS and counterstained with DAPI before mounting with coverslips.

Fluorescence images were acquired using a Zeiss Axio Observer microscope under standardized exposure conditions across all groups. Multiple fields encompassing the aortic valve region were imaged per section. Quantification of CD68-, 4HNE- and ACSL4-positive areas was performed using ImageJ by applying identical threshold settings across all images. Positive signal was expressed as percentage of total valve area. All image acquisition and analyses were performed by investigators blinded to treatment allocation.

For collagen assessment, Sirius Red staining was performed. Cryosections were air-dried, rehydrated through graded ethanol (100%, 90%, 70%), and rinsed in distilled water. Sections were stained with hematoxylin, washed under running water, and incubated in Sirius Red solution (0.1% Direct Red 80 in saturated picric acid) for 15 minutes. After washing, sections were dehydrated through graded ethanol, cleared in xylene, and mounted. Collagen content was assessed by light or polarized microscopy and quantified using ImageJ.

### Flow Cytometry

Peripheral immune cell populations were quantified by flow cytometry. Briefly, 100 µL of whole blood was collected from each mouse and processed immediately. Red blood cells were removed using red cell lysis buffer (BioLegend, #420301) followed by washing in phosphate-buffered saline. To prevent nonspecific antibody binding, Fc receptors were blocked using anti-mouse CD16/CD32 (Fc Block; Pharmingen) according to the manufacturer’s instructions. Cells were subsequently stained with fluorophore-conjugated antibodies against Ly6G, B220, CD11b (BioLegend), CD3 (eBioscience), CD45, and NK1.1 (BioLegend) for 20 minutes at 4°C in the dark. After washing, samples were analyzed on a FACSCanto II flow cytometer (BD Biosciences). Compensation was performed using single-stained controls, and data were analyzed using FlowJo software (Tree Star).

### Plasma Proteomics

Murine plasma samples were analyzed using the Olink Target 96 Mouse Exploratory panel according to the manufacturer’s instructions. Data were normalized and analyzed using OLINK NPX Signature software.

### Study Approval

The clinical study was approved by the Ethics Committee of the University of Bonn (approval number 077/14). All participants provided written informed consent. The study was conducted in accordance with the Declaration of Helsinki and Good Clinical Practice guidelines.

### Clinical Cohort

Patients with echocardiographic evidence of aortic valve sclerosis or mild-to-moderate AS were prospectively enrolled at the University Hospital Bonn, Germany. Severe AS and severe, life-threatening comorbidities limiting life expectancy were exclusion criteria. Baseline clinical characteristics were recorded at study entry. Patients underwent standardized transthoracic echocardiography at baseline and at 6-month intervals during follow-up. Peak transvalvular velocity, mean transvalvular gradient, and left ventricular ejection fraction were assessed in accordance with current guideline recommendations. Vitamin K status was assessed by measurement of circulating undercarboxylated osteocalcin (ucOC) using a commercially available ELISA kit (Takara, #MK118) according to the manufacturer’s instructions.

### Data Availability

The RNA-sequencing data generated in this study will be made publicly available upon publication in the Gene Expression Omnibus (GEO). The publicly available human aortic valve RNA-sequencing dataset re-analyzed in this study (non-diseased, fibrotic, and calcific tissue) was obtained from the Center for Interdisciplinary Cardiovascular Sciences (CICS) MultiOmics database, Brigham and Women’s Hospital, Harvard Medical School. All remaining data supporting the findings of this study are available within the article and its Supplemental Material, or from the corresponding author upon reasonable request.

### Statistical Analysis

Continuous variables are presented as mean ± SEM or as median (IQR), and categorical variables as counts and percentages. The distribution of continuous data was assessed for normality using the Shapiro-Wilk test, and parametric or non-parametric tests were applied accordingly. Two-group comparisons of normally distributed data were performed using the unpaired two-tailed Student’s t-test. Comparisons among more than two groups were performed using one-way ANOVA, and comparisons involving two factors using two-way ANOVA, each followed by Tukey’s multiple-comparisons test. Serial echocardiographic measurements were analyzed by two-way ANOVA with treatment and time as factors. In the clinical cohort, the association between log10-transformed ucOC and annualized progression (Δpeak velocity/year, Δmean gradient/year) was assessed by linear regression, with β coefficients and 95% CIs reported. An optimal ucOC cut-off (9.73 ng/mL) was derived by maximizing the Youden index. Stratified group comparisons used the unpaired two-tailed Student’s t-test. Multivariable linear regression with Δpeak velocity/year as the dependent variable adjusted for age, sex, baseline peak velocity, hypertension, diabetes, and hypercholesterolemia (Figure 6H). A prespecified sensitivity analysis was restricted to patients without CKD. A two-sided P < 0.05 was considered significant. Analyses were performed using GraphPad Prism (11.0.0) and IBM SPSS Statistics (31.0.0.0).

## Acknowledgements

The authors would like to thank Anna Flender and Rebecca Gunt for excellent technical assistance. Moreover, we thank Sandra Adler, Marta Stei, Gloria Martinac, and Pia Fuchs for assistance with coordination and technical assistance of wire injury surgeries. Parts of the figures were created with BioRender.

## Author Contributions

Conceptualization: ER and MAZ. Methodology: ER, AS, JO, GN, KC-N, MAZ. Investigation (experiments, data acquisition): ER, SS, AY, AS, DW, BB, RNJ, MB, MAZ. Formal analysis (statistics, bioinformatics): ER and MAZ. Data curation: ER, SS, AY, AS, DW, MAZ. Resources (patient samples, reagents, models): BA-K, JS, SB, MK, FB, GN, SZ. Visualization (figures): ER, SZ, MAZ. Writing – original draft: ER and MAZ. Writing – review & editing: all authors. Supervision: GN, SZ and MAZ. Project administration: ER and MAZ. Funding acquisition: ER, GN, SZ, MAZ. All authors read, critically revised, and approved the final manuscript.

---

**Table S1.**
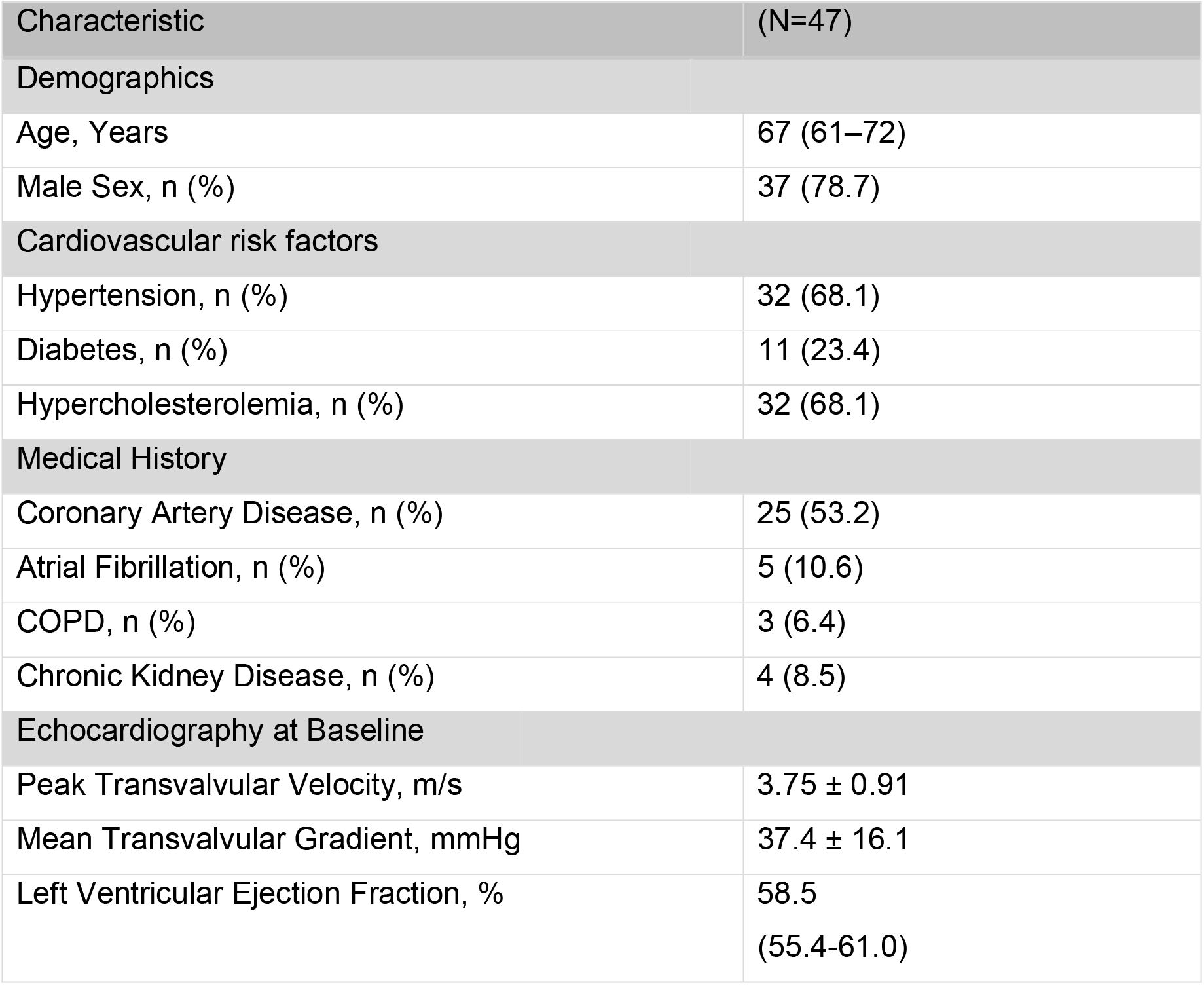
Baseline Characteristics Histology Cohor.

**Figure S1.**
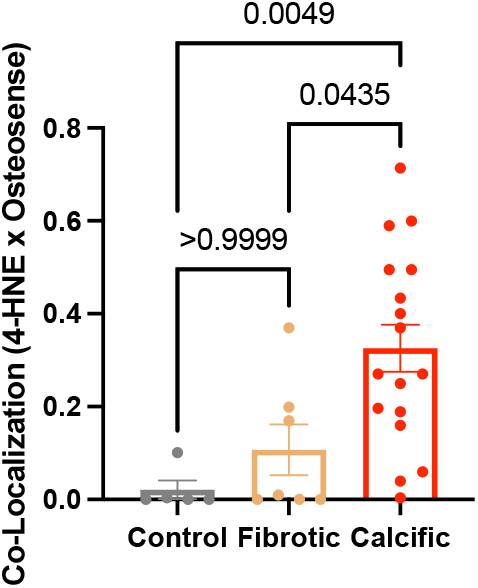
Lipid peroxidation co-localizes with calcification in human aortic valves. (A) Quantification of co-localization between 4-HNE (lipid peroxidation) and Osteosense680 (hydroxyapatite) in human aortic valve tissue from control, fibrotic, and calcific specimens. Data are presented as mean ± SEM. Statistical analysis was performed using one-way ANOVA with Tukey’s multiple-comparisons test.

**Figure S2.**
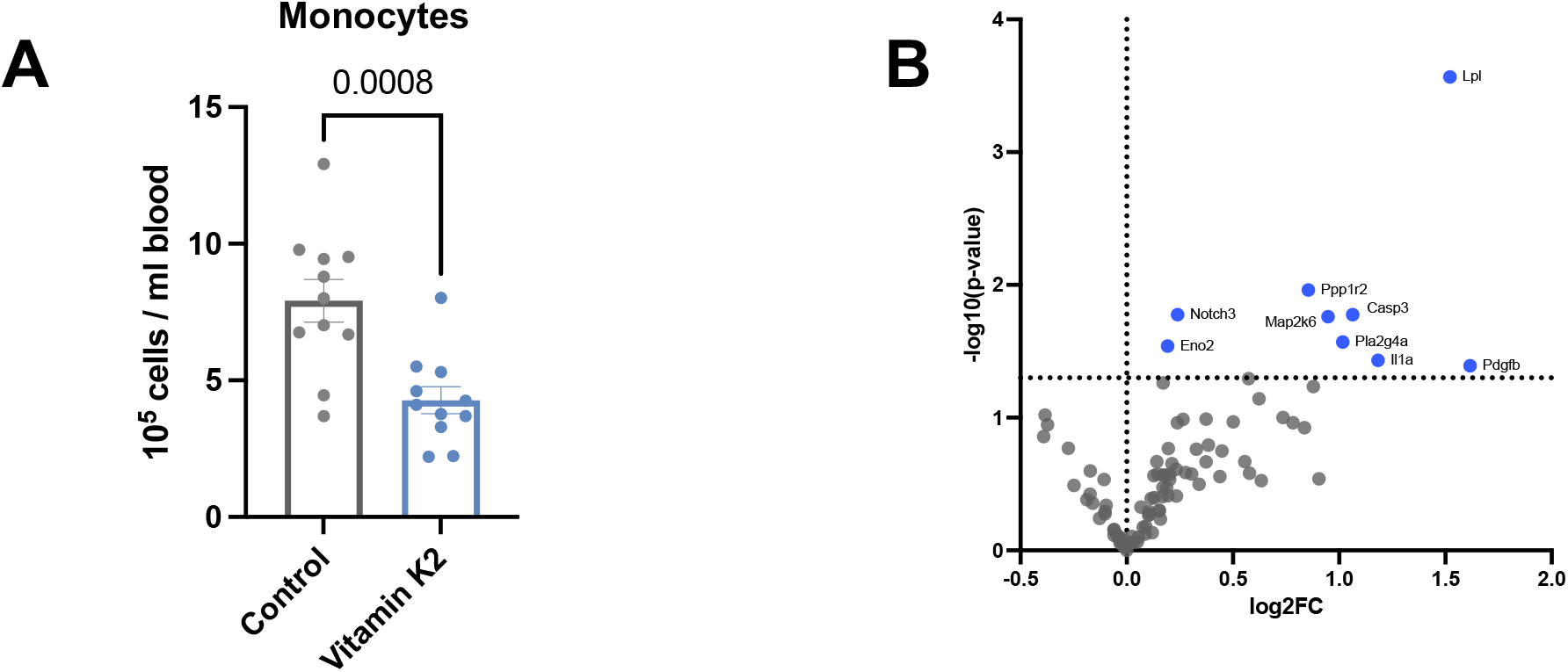
Vitamin K2 reduces circulating monocytes and modulates the plasma proteome in experimental aortic stenosis. (A) Quantification of peripheral blood monocytes by flow cytometry in control and vitamin K2–treated mice, expressed as 10⁵ cells per mL blood. (B) Volcano plot of plasma proteins measured by Olink proteomics in vitamin K2–treated versus control mice, showing log2 fold change against −log10(p-value). The vertical dotted line indicates no change (log2FC = 0) and the horizontal dotted line the significance threshold (p = 0.05); significantly regulated proteins are highlighted in blue and labeled. Data in (A) are presented as mean ± SEM and were analyzed using an unpaired two-tailed Student’s t-test.

